# An Efficient Moments-Based Inference Method for Within-Host Bacterial Infection Dynamics

**DOI:** 10.1101/116319

**Authors:** David J. Price, Alexandre Breuzé, Richard Dybowski, Piero Mastroeni, Olivier Restif

## Abstract

Over the last ten years, isogenic tagging (IT) has revolutionised the study of bacterial infection dynamics in laboratory animal models. However, quantitative analysis of IT data has been hindered by the piecemeal development of relevant statistical models. The most promising approach relies on stochastic Markovian models of bacterial population dynamics within and among organs. Here we present an efficient numerical method to fit such stochastic dynamic models to *in vivo* experimental IT data. A common approach to statistical inference with stochastic dynamic models relies on producing large numbers of simulations, but this remains a slow and inefficient method for all but simple problems, especially when tracking bacteria in multiple locations simultaneously. Instead, we derive and solve the systems of ordinary differential equations for the two lower-order moments of the stochastic variables (mean, variance and covariance). For any given model structure, and assuming linear dynamic rates, we demonstrate how the model parameters can be efficiently and accurately estimated by divergence minimisation. We then apply our method to an experimental dataset and compare the estimates and goodness-of-fit to those obtained by maximum likelihood estimation. While both sets of parameter estimates had overlapping confidence regions, the new method produced lower values for the division and death rates of bacteria: these improved the goodness-of-fit at the second time point at the expense of that of the first time point. This flexible framework can easily be applied to a range of experimental systems. Its computational efficiency paves the way for model comparison and optimal experimental design.

**Author Summary:** Recent advancements in technology have meant that microbiologists are producing vast amounts of experimental data. However, statistical methods by which we can analyse that data, draw informative inference, and test relevant hypotheses, are much needed. Here, we present a new, efficient inference tool for estimating parameters of stochastic models, with a particular focus on models of within-host bacterial dynamics. The method relies on matching the two lower-order moments of the experimental data (i.e., mean, variance and covariance), to the moments from the mathematical model. The method is verified, and particular choices justified, through a number of simulation studies. We then use this method to estimate models that have been previously estimated using a “gold-standard” maximum likelihood procedure.

**List of symbols:**

- *A*: number of animals
- *T*: number of tagged strains
- *n*: number of organs
- *N_i_*: number of bacteria in organ *i*
- *m_ij_*: migration rate from organ *i* to organ *j*
- *k_i_*: killing rate in organ *i*
- *r_i_*: replication rate in organ *i*
- *τ_i_*: observation time *i*
- **A**, **B**, **C**: matrices
- ***λ***: vector of transition rates
- *B*: Number of bootstrap samples
- ***θ**^∗^*: MDE parameter estimate

**Abbreviations:**

- ABC: approximate Bayesian computation
- IT: isogenic tagging
- LV: live vaccine
- MARE: mean absolute relative error
- MDE: minimum divergence estimate
- MLE: maximum likelihood estimate
- qPCR: quantitative polymerase chain reaction
- WITS: wildtype isogenic tagged strain

## 1 Introduction

The elucidation of basic kinetic rates governing bacterial growth during infection (such as division and death rates) has been recognised as an important challenge for over 60 years [1]. Thanks to recent technological developments in microbiology, and pushed by growing concern over antimicrobial resistance and the need for new vaccines, the last decade has witnessed rapid progress in the quantification of *in vivo* dynamics of bacterial infection in animal models. Two experimental approaches in particular have shown great promise across multiple pathogen species: isogenic tagging, the focus of this report, and fluorescence dilution, a term encompassing several techniques from which bacterial replication can be inferred [2]. Isogenic tagging (IT) consists in generating an arbitrary number of sub-clones of a given bacterial strain, each defined by a unique genetic tag (a predetermined nucleotide sequence) inserted in a non-coding region of the chromosome. When grown together *in vitro* or *in vivo*, every tagged strain behaves identically to the original strain. Their relative frequencies within a bacterial culture can be measured by quantitative qPCR or sequencing of the tagged region. Taken together with the absolute number of viable bacteria (e.g., by plating colonies), changes in the frequencies of the tags within the bacterial population can reveal underlying variations in the rates at which bacteria divide, die and disperse. For example, a constant number of viable bacteria accompanied by a loss of some of the tags in a closed population would indicate that a certain proportion of bacteria have died and been replaced by replication. Likewise, when monitoring tag frequencies in two or more anatomical compartments within animals, a gradual homogenisation among organs can reveal the transfer of bacteria. While some studies have stopped at qualitative interpretations of such empirical patterns [3, 4, 5, 6], it is possible to quantify underlying processes with the help of mathematical models. Two different types have been used: stochastic population dynamic models to estimate bacterial division, death and migration rates [7, 8, 9, 10, 11, 12], and population genetic models to estimate bottleneck sizes [13, 14]. Our aim is to develop efficient inference methods to deal with the former type of models.

Stochastic birth–death–migration models (a canonical class of Markovian processes [15]) are a common choice to analyse IT experiments, and naturally lead to likelihood–based inference, using either maximum likelihood [9, 11] or Bayesian estimation [12]. In a given experiment, assuming that all of the *A* animals sampled at a given time and in given conditions are identical, and that all of the *T* tagged strains infecting each animal act independently of each other and are governed by identical rates, we can treat the *A × T* observed strain abundances as independent realisations of a stochastic birth–death–migration process, and calculate the likelihood of any model of interest accordingly. In most published IT studies, bacteria can grow at different rates in different locations within an animal, and migrate from one location to another, generating a network of subpopulations (or metapopulation). This increases the dimensions of both the state variable space (as the model must keep track of the multivariate distribution of bacterial abundance) and the parameter space. Calculating the likelihood of such a model given an experimental dataset requires solving a complex stochastic model, which will rarely be possible analytically. Even a linear birth–death process with a non–Poisson immigration process (representing the transfer of a finite inoculum dose) is sufficient to prevent a fully analytical treatment, and results in a computationally intensive estimation process [12]. Alternatively, approximate–likelihood (e.g., iterative filtering [16]) or likelihood–free (e.g., approximate Bayesian computation or ABC [17]) methods involve the generation of a large number of stochastic simulations of the model of interest, which can be equally time–consuming, even when taking advantage of parallel computation. Although this may not be a problem when fitting a single model to a single dataset, it limits our ability to compare multiple models across complex datasets (typically involving multiple experimental treatments) and, beyond that, use these inference tools for the purpose of optimising experimental design [18]. Hence, there is a need for alternative inference methods using suitable approximations to achieve greater gains in computational efficiency.

The dynamics of multivariate Markovian processes can be approximated using moment– closure methods [19]. Mathematically, a system of differential equations for the moments of the state variables can be derived analytically from the governing equation of any stochastic model [20]. By effectively ignoring the higher moments, a closed, small-dimension system can be derived, allowing fast numerical solution of the lower moments at any time point. Parameter estimation can then be achieved by fitting the first and second order moments of the model to the mean, variance and covariance of the corresponding variables in the data. Apart from a few proof–of–principle studies using simulated chemical reaction data [21, 22, 23] that show great promise, application to statistical inference from biological data remain scarce. As a rare example, Buchholz et al. [24] implemented a moment–based method to solve a multiple T–cell differentiation pathway problem, fitting the moments of a large number of alternative stochastic models to experimental data using a *χ*^2^ statistic. This suggests that efficient moment–based inference methods should be made more readily available to unleash the full potential of stochastic models in experimental biology.

Our objective is to provide a functional and flexible computational framework to estimate the parameters of stochastic metapopulation models for the within–host dynamics of infection, and demonstrate its application and value to analyse IT studies. The model tracks the probability distribution of the number of copies of a tagged strain of bacteria across a network of anatomical compartments within an animal. The goal is to estimate the bacterial division and death rates within each organ, and migration rates between each pair of compartments. First, we present an algorithm that evaluates the first- and second-order moments of the state variables for arbitrary network structures, and assess its accuracy and speed against a gold standard for stochastic models: the exact Gillespie algorithm. We then compare the accuracy of several inference options against simulated data, and finally apply the most promising method to a likelihood-based method in order to assess the quality of inference and increased computational efficiency. The massive gain in speed compared to likelihood-based inference allows us to use parametric bootstrap to quantify parameter uncertainty and goodness-of-fit. Finally, we provide a re-analysis of a recent dataset on the dynamics of *Salmonella enterica* serovar Typhimurium in the blood, liver and spleen of vaccinated mice [11]. We also demonstrate how empirically derived noise terms (e.g., caused by imprecise data collection) can be taken into account.

## 2 Methodology

### 2.1 Biological context

We consider the general case of a bacterial pathogen inoculated into an animal host where it can potentially reach *n* anatomical compartments—which can be distinct organs, tissues, lumens, or predefined sections thereof. All our examples are motivated by IT experiments in which a set of identical animals receive the same initial inoculum dose in one compartment (e.g., mouth, nose, blood, peritoneum, etc) at time *t* = 0. The inoculum is composed of an even mix of *T* tagged strains. At given times *τ*_1_,*τ*_2_ etc, a subset of *A*_1_, *A*_2_, etc animals are chosen at random and euthanised. The abundance of each tagged strain in each of the *n* anatomical compartments of interest is measured. Thus, at a given time *τ_i_*, the data consist of a matrix **D**_*i*_ with *n* rows by *A_i_T* columns, filled with observed bacterial numbers. From this matrix, we can calculate the observed moments, namely the mean and variance of strain abundance within each compartment, and the covariance between each pair of compartments.

Depending on the experimental procedures, these observations are usually subject to some degree of uncertainty, due to observational error. In general we assume that this error is random with a mean of zero, so that there is no systematic bias; this should be assessed by the researchers who conducted the experiments. As a result, we assume that the observed means are unbiased, but the observed (co)variances may be incorrect. In Section 2.7, we describe how known sources of error can be accounted for as part of the data processing procedure. In addition, there usually is some uncertainty about the actual inoculum dose received by each animal. Variations in the abundance of each strain should be assessed experimentally by testing several inoculum doses: this provides estimates for the initial mean and variance of the number of bacteria with each tag that are present in the target compartment at *t* = 0.

We emphasise a few key assumptions and caveats of the present study, which we review in further detail in the Discussion. First, the variable of interest from a modelling perspective is the abundance of a single tagged strain, rather than the total bacterial load per animal (as the latter can be obtained by adding up individual strains). Indeed, our model framework assumes that, over the time period considered and for a given set of initial conditions, the rates of bacterial division, death and migration *per capita* are independent of the total bacterial load. Second, we assume that all the bacterial cells are governed by identical rates of division, death and migration. While this excludes the case of so–called persister cells (i.e., a subset of bacteria with a much lower division rate than the rest) or similar discrete partition, it is worth noting that the model we describe below does generate stochastic variations in the generation time of bacteria, consistent with empirical distributions of bacterial replication in vivo [25].

### 2.2 Stochastic model framework

As a function of time *t* since inoculation, the vector of positive integer state variables ***N*** (*t*) = {*N*_1_(*t*), *…, N_n_*(*t*)} represents the simultaneous abundance of bacteria in compartments 1 to *n*. In the context of IT studies, this represents a single tagged strain. Three types of stochastic events drive the bacterial dynamics: division (which adds one bacterium to a given compartment), death (which removes one bacterium from a given compartment) and migration (which moves one bacterium from one compartment to another). Assuming linear transition rates, we have a total of *n* division rates *r_i_N_i_* and death rates *k_i_N_i_* within organ *i*, and *n*(*n −* 1) migration rates *m_i,j_ N_i_* from compartment *i* to *j*. Note that specific models may assume that some of the parameters are equal to zero, for example if there is no physical connection between given pairs of compartments.

In particular, we consider two geometries, illustrated in Figure 1, corresponding to two typical anatomical topologies of relevance to bacterial infection: a radial network with a central compartment (e.g., bloodstream supplying every organ), and a linear network (e.g., digestive track).

**Figure 1.**
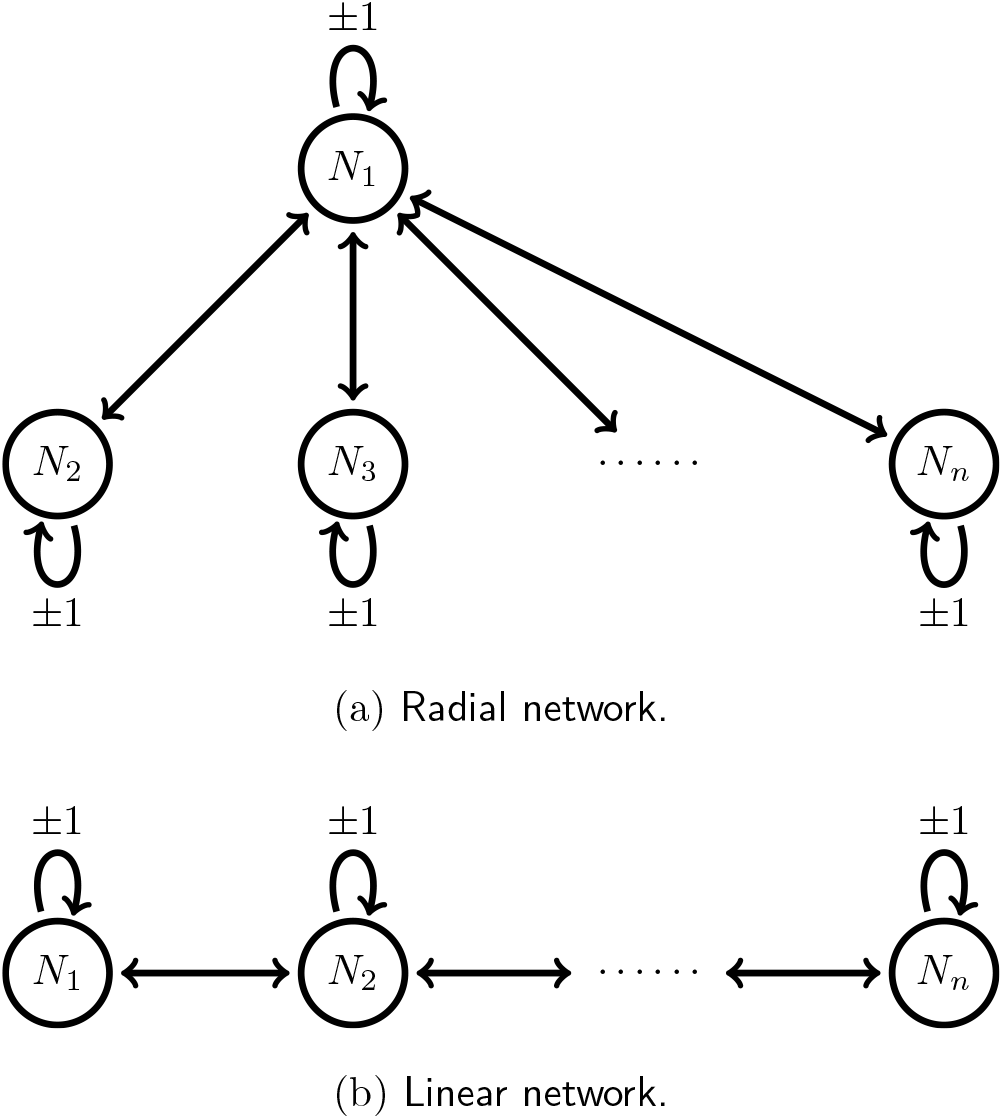
Diagram illustrating the two types of network structure we consider.

### 2.3 Computation of the first-and second-order moments

The method we propose for parameter inference relies on the first two moments of the stochastic system. That is, we use only the expected number of bacteria within each compartment, the variance of the number of bacteria within each compartment, and the pair-wise covariances. A simple approach to generating these moments for a particular stochastic system in terms of the model parameters, is given by [26]. Letting ***λ*** = {*r*_1_*N*_1_, *…, r_n_N_n_, k*_1_*N*_1_, *…, k_n_N_n_, m*_1,2_*N*_1_, *…*} be the vector of transition rates, we can write the *h^th^* non-central moment of the state of the *i^th^* compartment as:

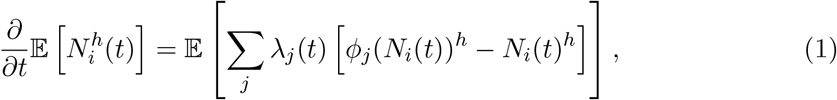

where *φ_j_* is a function describing the change of the state for the *j*th transition. In Supplementary Materials (S1.7), we show that this leads to a closed, linear system of differential equations for the first moments. Letting ***M***_1_(*t*) = {𝔼 [*N_i_*(*t*)], 1 *≤ i ≤ n*} be the vector of first moments as a function of time, we can express these differential equations in matrix form as:

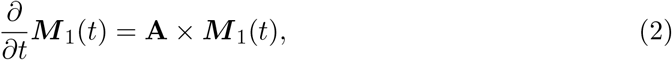

which leads to the solution,

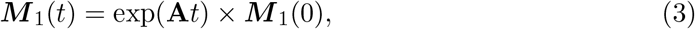

where ***M***_1_(0) are the initial conditions of the system, **A** is a time-independent matrix containing the model parameters, and *exp* is the matrix exponential function.

Next, let ***M***_2_(*t*) = {𝔼 [*N_i_*(*t*)*N_j_* (*t*)], 1 *≤ i ≤ n*, 1 *≤ j ≤ n*} be the vector of second moments. By applying Duhamel’s formula to the differential equations obtained from equation (1), we obtain the following expression for the second-order moments:

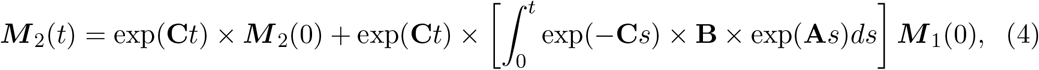

where **B** and **C** are time-independent matrices containing model parameters, and ***M***_1_(0) and ***M***_2_(0) are vectors containing the initial moments. Using the numerical method for matrix exponential in [27], we can evaluate the first- and second-order moments at any time point. Remarkably, no moment-closure approach is required as the expressions for the second-order moments are independent of higher-order moments. See Supplementary Information for a full derivation.

### 2.4 Parameter Inference by Divergence Minimisation

Given a dataset consisting of one or more matrices **D** of bacterial counts (as per section 2.1), and a stochastic model (as per section 2.2), we now describe methods to estimate the parameter values of the model that minimise the divergence between the predicted and observed distributions of bacterial abundance, using only the lower moments of those distributions. Specifically, we evaluate the means, variances and covariances of the *N_i_* variables at a given time *t*. From the corresponding matrix **D**, we calculate the vector of observed means ***µ***^(**D**)^ and the matrix of observed variance-covariance **V**^(**D**)^; and from the model’s solution given a set of parameters ***θ***, we compute the vector of predicted means ***µ***^(*θ*)^ and the matrix of predicted variances-covariances **V**^(*θ*)^ which can be derived from ***M***_2_(*t*). We compared four common divergence measures: a Chi-Squared metric, and normal approximations to the Mahalanobis distance, the Hellinger distance, and the Kullback-Leibler divergence.

Note that none of the measures below is designed to deal with the particular situation when any of the organs is reported void of bacteria (i.e., *N_i_* = 0) in all replicates at a given time point, i.e. if all the observed moments related to that organ are equal to zero. In some experimental systems, this may occur as an artefact of the observation method, e.g., when only a small sample is measured: in this case, it is possible to “correct” the data for sampling biases (see Section 2.7). Otherwise, a simple solution would be to remove the moments relative to that organ from the inference procedure. In some cases, it may make sense to completely remove the empty organ from the model if no meaningful inference can be expected from its inclusion, as illustrated in Section 3.5.

The Chi-Squared metric adds up the squared pairwise-differences between each predicted moment and its corresponding observed moment, each term being scaled by the magnitude of the observed moment. As a result, all moments are effectively treated equally. The expression for this divergence is:

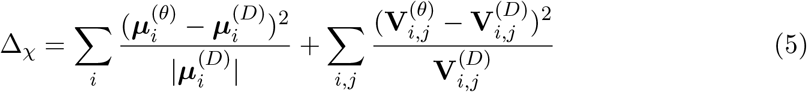

The other three divergence expressions we tested make use of the fact that our chosen statistics ***µ*** and **V** are moments of distributions. Since we do not establish complete characterisations of the distributions, we decided to borrow the expressions of well-known divergences measures for multivariate normal distributions, as these only require the knowledge of their first and second-order moments. In other words, we compute the divergence between two multivariate normal distributions with respective moments (***µ***^(**D**)^, **V**^(**D**)^) and (***µ***^(***θ***)^, **V**^(***θ***)^).

The Mahalanobis distance is measured between each point in the observed data, and the distribution described by the predicted moments. That is, by estimating the parameters we are trying to find the distribution that these data are most likely to have been sampled from. Specifically, each observation ***X_j_*** is given by column *j* of the data matrix **D**. The divergence is obtained by summing the distances to the predicted distribution from every observation:

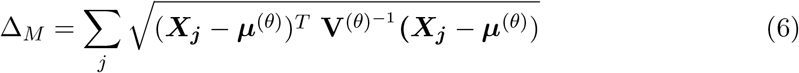

The squared Hellinger distance is given by:

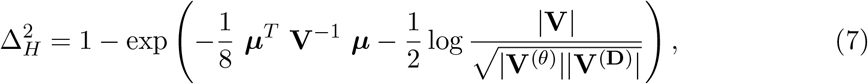

where ***µ*** = ***µ***^(*θ*)^ *− **µ***^(**D**)^ and 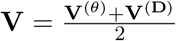.

The Kullback-Leibler divergence from the predicted distribution to the observed one is given by:

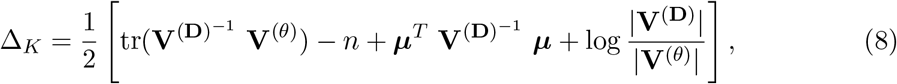

where tr(*·*) is the trace operator. The derivations of the Hellinger and Kullback-Leibler divergences are given in Supplementary Materials S1.2.

Even though these expressions may not provide correct estimates of the actual “Mahalanobis distance”, “Hellinger distance” and “Kullback-Leibler divergence” between the data and the predicted distributions (as these are not normally distributed), they still provide adequate divergence measures: they all return positive values which are only equal to zero (hence are minimised) when ***µ***^(*θ*)^ = ***µ***^(**D**)^ and **V**^(*θ*)^ = **V**^(**D**)^, which occurs when the data are drawn from the predicted distribution. Furthermore, we only require that the calculated divergence here is ordered in the same way as the true divergence between these distributions, so that the minimum occurs at the same parameter values. We use these properties for the purpose of parameter inference, given a dataset and a model. The set of parameter values ***θ*** that minimises the divergence measure is termed the “minimum divergence estimate” (MDE).

All code was written in R [28]. We used the UObyQA optimisation routine in the powell package [29].

### 2.5 Bootstrap Variance Estimate

In order to quantify the uncertainty in our parameter estimates conditional on the dynamic model considered, we utilise the parametric bootstrap method (e.g., [30]), which can be exploited simply here due to the computational efficiency of the inference approach. Having obtained an MDE, ***θ**^∗^* for a given data matrix **D** and a given model, we simulate *B* data sets from the model at these parameters (i.e., ***x**^b^ ∼ f* (*x | **θ**^∗^*), *b* = 1, *…, B*). For each simulation, we estimate the corresponding parameters using the MDE technique (***θ**^b^*, *b* = 1, *…, B*). These *B* estimated parameters are used to estimate the variance-covariance matrix of the parameter estimates. Subsequently, one can use this matrix to estimate confidence intervals in a number of ways [31]. For simplicity, the analysis in Section 3.5 uses the ellipse package in R [32] to obtain approximate 95% confidence intervals of the model parameters (that is, assuming the bootstrapped parameter estimates follow a multivariate normal distribution).

### 2.6 Model Goodness-of-fit

In order to assess the model goodness-of-fit, we once again utilise the parametric bootstrap. Concurrent to calculating the uncertainty estimates using the MDE method, the divergences corresponding to each simulated data set at their estimated parameter values are recorded. These bootstrapped divergences can thus be used to represent the null distribution of divergences for the model at the estimated parameter values – giving a representation of the divergences we should expect from the model at these values. The divergence estimated for the observed data is then compared to the null distribution to obtain a p-value for the hypothesis that the data could have been generated by the model.

### 2.7 Data analysis with observational noise

A common source of error when fitting a model to experimental data is the observation process. In the type of microbiology experiments considered here, where the data represent bacterial loads in infected animals, there are at least two steps that affect the accuracy of the measurements: sampling (when only part of the bacterial loads are recovered) and and quantitation (the process by which the number of bacterial cells in the samples is measured). For example, in a recent IT experiment [11], sampling error was modelled as a binomial process (as known fractions of each homogenised organ were plated) and quantitation error was modelled using a log-normal distribution which was estimated empirically using an independent control experiment (in which known numbers of bacterial colonies were processed by qPCR in the same way as bacterial samples extracted from animals in the main experiment). Although both error distributions were centred on the true bacterial numbers (i.e. the mean numbers of bacteria were not biased), the variance and covariance in the reported data would not have been accurate estimates of the variance and covariance within the animals: hence our MDE could be biased if we did not account for observational errors.

In our present reanalysis of those data, we integrate both error terms before performing parameter inference, using the following procedure. Our goal is to propose a simple heuristic which could be applied to any experimental dataset with known (or assumed) observational error distributions. First, we generate a large number of stochastic simulations of the model under a biologically reasonable range of parameter values (i.e. using uniform prior distributions across sensible ranges) to generate “perfect” observations (1. in Figure 2). We then calculate the corresponding “perfect moments” from every simulated dataset (1M. in Figure 2), which we cannot observe from our experimental procedure. Next, we apply the observation noise to the simulated data (2. in Figure 2), as is the case in the experimental procedure, and calculate the corresponding “observed moments” (2M. in Figure 2). We then use linear regression models to establish a relationship between each of the perfect and observed moments (with transformations where appropriate), with weights given by the simulations proximity to the actual observed moments from the experiment. This calibrated regression model is then applied to the moments of the experimental data of interest, in order to estimate the moments of the true, unobserved bacterial loads in the animals. We eventually compute MDE using these corrected moments.

**Figure 2.**
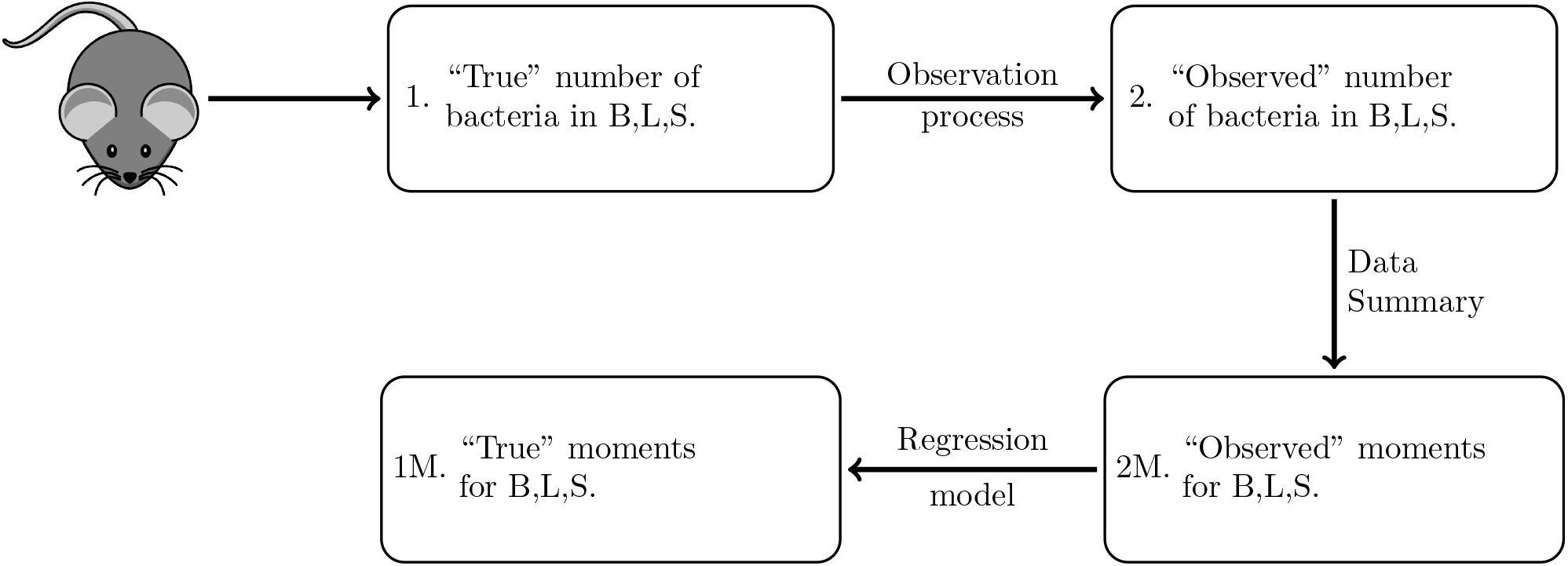
Schematic demonstrating the experimental procedure to obtain the observed moments. Ideally, we want to observe the “true” moments, however this is not possible due to the observation process. In order to understand the effect of the observation noise on the moments, we simulate the system, add noise consistent with the experimental system, and evaluate the relationship between the “true” and “observed” moments of these simulations. We can thus infer the “true” moments of the system, given our “observed” moments.

We note that for some models of the observation process, it may be possible to establish analytic relationships between the “perfect” and “observed” moments using conditional expectation and variance theory (e.g., [33]). The choice of suitable correction methods for a given system will depend on both the model complexity and the level of empirical quantification of observation noise.

## 3 Results

We begin by demonstrating the speed and accuracy of the computation of the first- and second-order moments for arbitrary network structures, compared to Gillespie simulations. We then assess the four previously mentioned divergence measures to validate our minimum divergence estimation procedure with each model, across a range of parameter values. Finally, we apply our inference method to reanalyse a published experimental dataset, and compare the results with the previous maximum likelihood estimates.

### 3.1 Moment computation

As a proof-of-concept, we compare our proposed computation method of the means, variances and covariances of bacterial loads, with the values derived from large numbers of Gillespie simulations, across a range of model structures and parameter values. Our results illustrate the computational effort required to obtain the same level of accuracy with each method. For each model and each parameter set, we simulated 100 experiments, each consisting of 10, 50, 100, 250, 500, or 1000 replicates (representing the product of the number of animals *A* by the number of tagged strains *T* as per section 2.1) at a given time, without observational error. The initial bacterial loads at *t* = 0 in each replicate experiment were drawn at random from a Poisson distribution, to mimic typical variability in inoculum doses in experiments [11]. We considered three model structures, illustrated in Fig. 3: (a) basic migration-birth-death model, (b) four-compartment radial network, and (c) four-compartment linear network. For each virtual experiment, we calculated the first-and second-order moments of the bacterial loads in two ways at a given time point (*t* = 6 for the basic migration-birth-death model, and *t* = 4 for the four-compartment radial and linear network).

**Figure 3.**
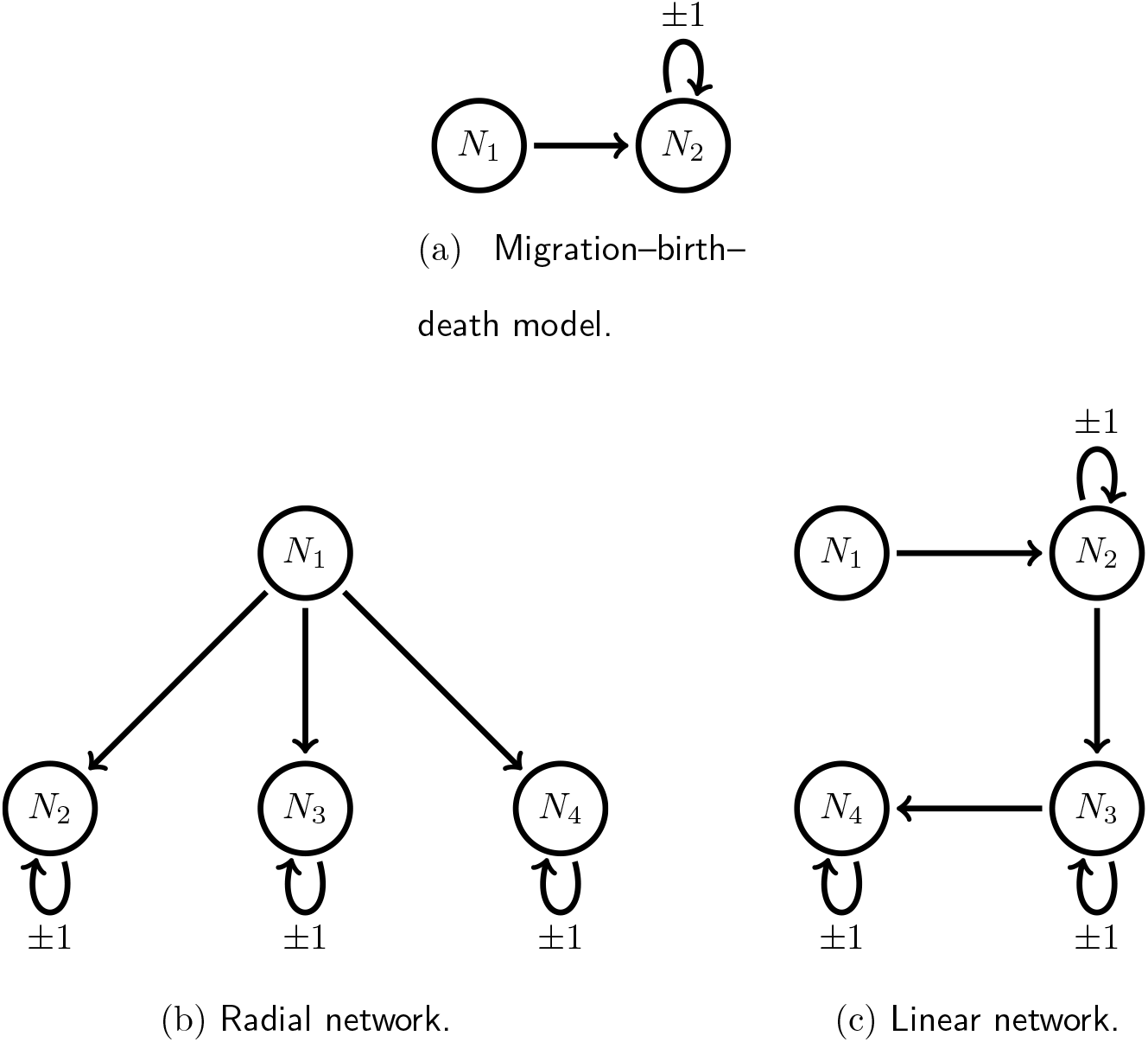
Illustrations of three representative model structures.

We randomly drew the stated number of initial conditions (ranging from 10 to 1000), and evaluated the moments at the future time point in the following two ways. First, we used Gillespie simulations to progress each initial condition forward to the stated observation time, and then calculated the moments from the collection of simulations. Second, we took the stated number of initial conditions to estimate the moments at time zero, and used these to evaluate the moments at the observation time using equations (3) and (4); we refer to this as the direct moment-calculation method.

Results are included in the Supplementary Materials. Figure S1 shows that there is still a greater amount of variation in the moments from 1000 Gillespie simulations, compared to the direct approach. Both methods produce more accurate estimates of the moments as experiment size increases, as a direct result of more accurate estimates of the initial distribution of bacteria. Furthermore, the computation time of the Gillespie approach increases exponentially with the number of simulations, while the direct method is consistently more efficient, independent of the size of the experiment (Figure S2). Similar patterns are observed for the four-compartment linear (Figure S3) and radial (Figure S4) network models.

### 3.2 Divergence Measures: Simulation Study

Next, we compare the accuracy of MDE among the four candidate divergence measures, using data simulated from the same three model structures as in the previous section and across a range of parameter values. For each model structure and each parameter combination, we ran 100 series of 100 Gillespie simulations from *t* = 0 to each of 8 observation times, representing 800 experiments (each of size 100) with “perfect” observations of the system (i.e., without experimental noise). This process was repeated for a set of different parameter values for each model to represent different scenarios – three representative values for each parameter in the basic model (3^3^ = 27 scenarios), and five randomly generated sets of parameters for the four compartment linear and radial networks with parameters. The initial conditions for each simulation, in every scenario, were randomly generated from a Poisson distribution with mean parameter 200. The optimisation routine was initiated at randomly generated conditions each time, with each parameter values drawn independently from a Uniform(0,1) distribution.

From the results of each simulated experiment, we computed the MDE using each of the Chi-Squared, Mahalanobis, Hellinger, and Kullback-Leibler divergences. The performance of each divergence was measured by the mean absolute relative error (MARE). That is, under scenario *s*, with *p* target parameters *θ_s_* = (*θ_1s_*, *θ_2s_*, …, *θ_ps_*), and estimated parameters *θ̂_sj_* = (*θ̂_1sj_*, *θ̂_2sj_*, …, *θ̂_psj_*) for the *j^th^* simulation, the MARE is given by:

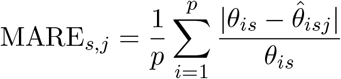

Figure 4 displays the average error across each scenario for the four divergence measures, at a range of observation times, for each of the three models under consideration. We can see that both the Hellinger and Kullback-Leibler divergence measures perform considerably better than the Chi-Squared and Mahalanobis divergences for the Basic, Linear and Radial networks. The similar performance of the Hellinger and Kullback-Leibler divergences is not unexpected, due to their relationship. We observe a marginal advantage in favour of the Kullback-Leibler divergence for the basic and radial network, and thus we use the Kullback-Leibler divergence in analysing experimental data in Section 3.5. The results aggregated by scenario for each model are shown in Supplementary Materials S1.4.

**Figure 4.**
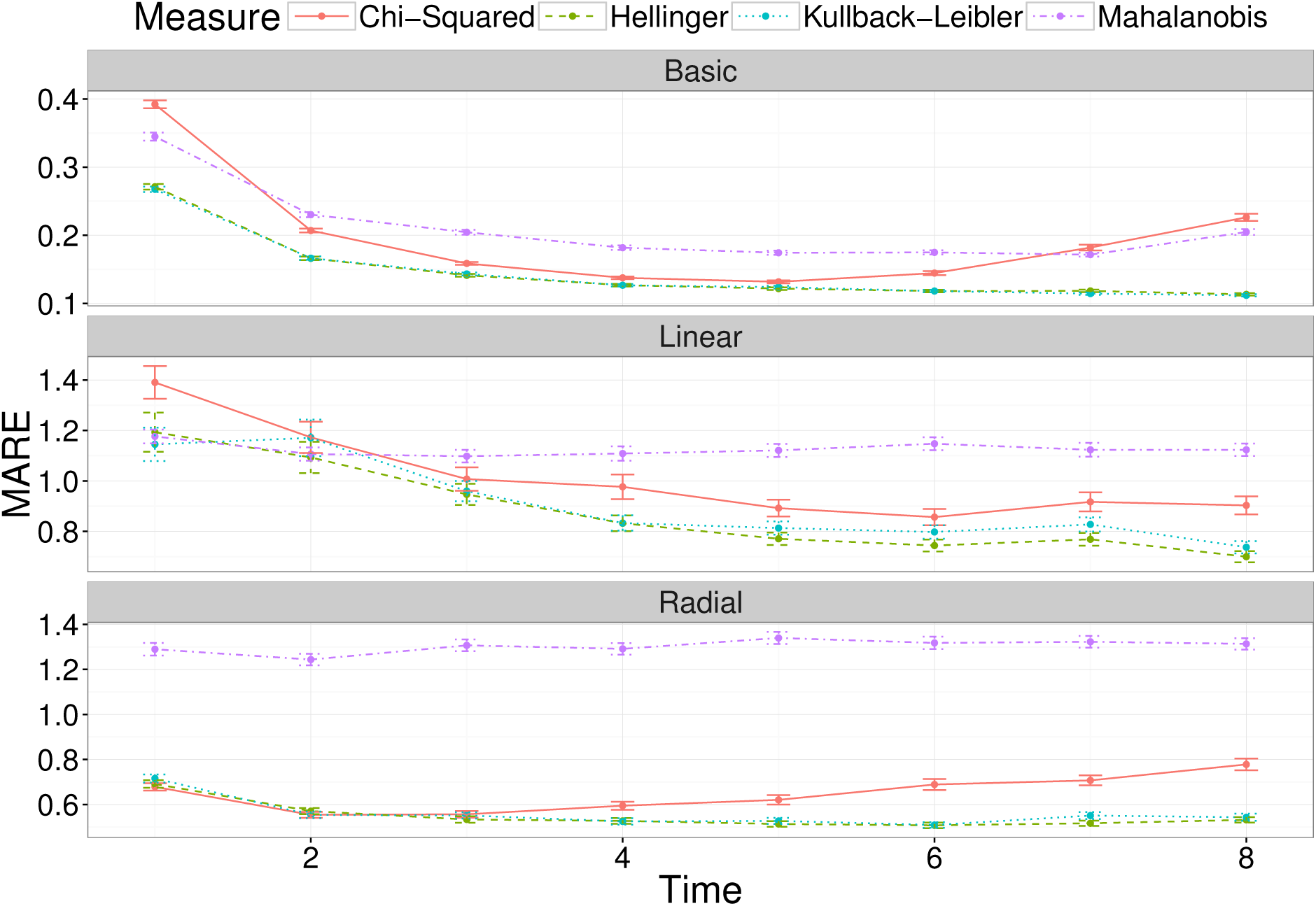
The average mean absolute relative error (MARE) across a number of observation times, for the two-compartment network (top panel), the linear four-compartment network (middle panel) and the radial four-compartment network (bottom panel). Each point is the average for that measure, at that observation time, with error bars representing the standard error.

Figure 5 further illustrates the distribution of parameter estimates for a few selected scenarios and time points (note that Scenario’s 4 and 12 just correspond to the combination of values for the parameters *m*_12_, *k*_2_, *r*_2_). For each time point, the variation was generated by the stochastic birth-death-migration process, demonstrating the level of uncertainty in parameter estimates due to the biological process itself. In this case, each simulated experiment was made up of 100 simulations; fewer replicates would increase the parameter uncertainty. At each time point, the estimates of the migration rate *m*_12_ are much less variable than those of the killing rate *k*_2_ and replication rate *r*_2_: this is actually caused by a very strong positive correlation between the latter two parameters (Figure 6).

**Figure 5.**
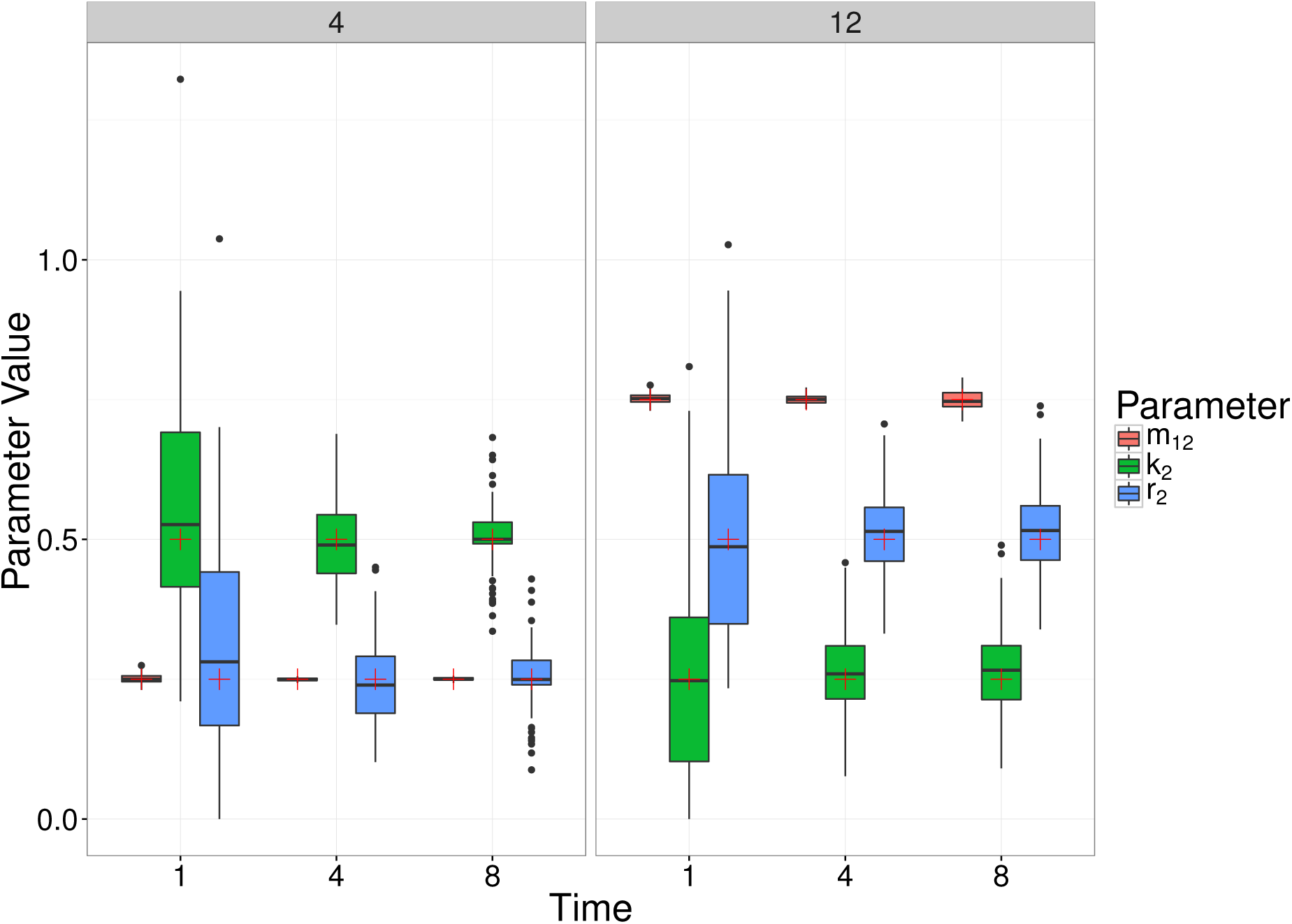
Box plots of MDE’s corresponding to simulated data for two different choices of parameter values (Scenario 4 and 12), at a number of different observation times (1, 4 and 8) of the simple, migration birth-death model. The box plots represent 100 parameter estimates corresponding to the 100 simulations, and the red crosses denote the true parameter value.

**Figure 6.**
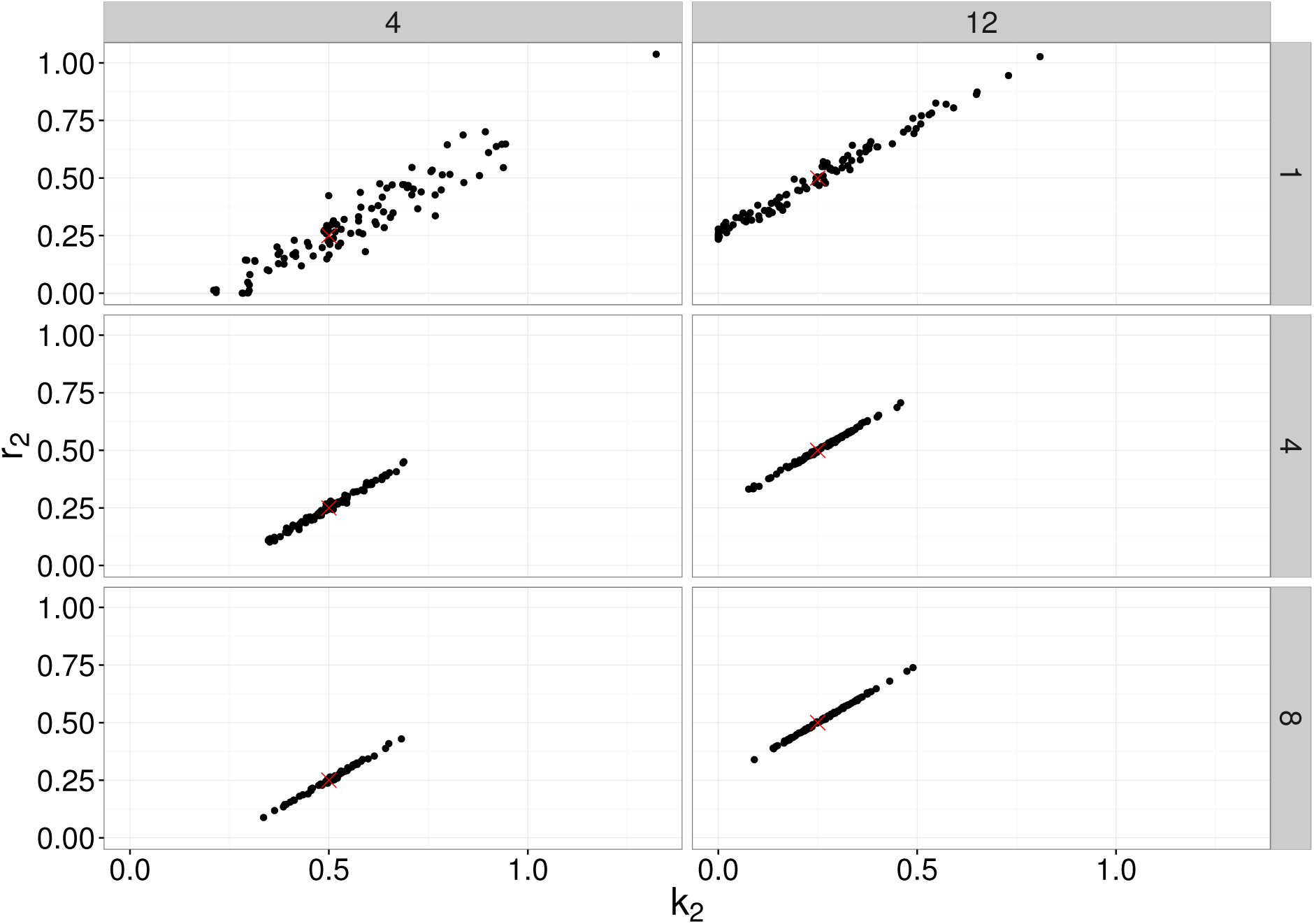
Bivariate plot of MDE’s corresponding to simulated data for two different choices of parameter values (Scenario 4 and 12), at a number of different observation times (1, 4 and 8) of the simple, migration birth-death model. The points represent 100 parameter estimates of *r*_2_ and *k*_2_ corresponding to the 100 simulations, and the red crosses denote the true parameter value.

This can be understood intuitively, as the net growth rate *r*_2_ *− k*_2_ and the migration rate *m*_12_ are the main determinants of the mean bacterial load in compartment 2. We also remark that the earlier observation time typically contains more information about the migration rate, whereas the later observation time better captures the replication and killing rates. This is a result of the particular dynamic we are considering, whereby bacteria begin in the first compartment, and steadily move out to the other compartment(s). For a relatively large migration rate, the first compartment is typically close to depleted at later observation times, and thus we obtain a poorer estimate (e.g., Scenario 12, Figure 5). For a smaller migration rate, there will be an optimal observation time that has allowed sufficient numbers of bacteria to have moved out of the first compartment, improving the parameter estimate (e.g., Scenario 4, Figure 5). Results of all scenarios are shown in Supplementary Materials S1.6. Figures 7 and 8 show similar results to Figure 5, for the linear and radial examples, respectively.

**Figure 7.**
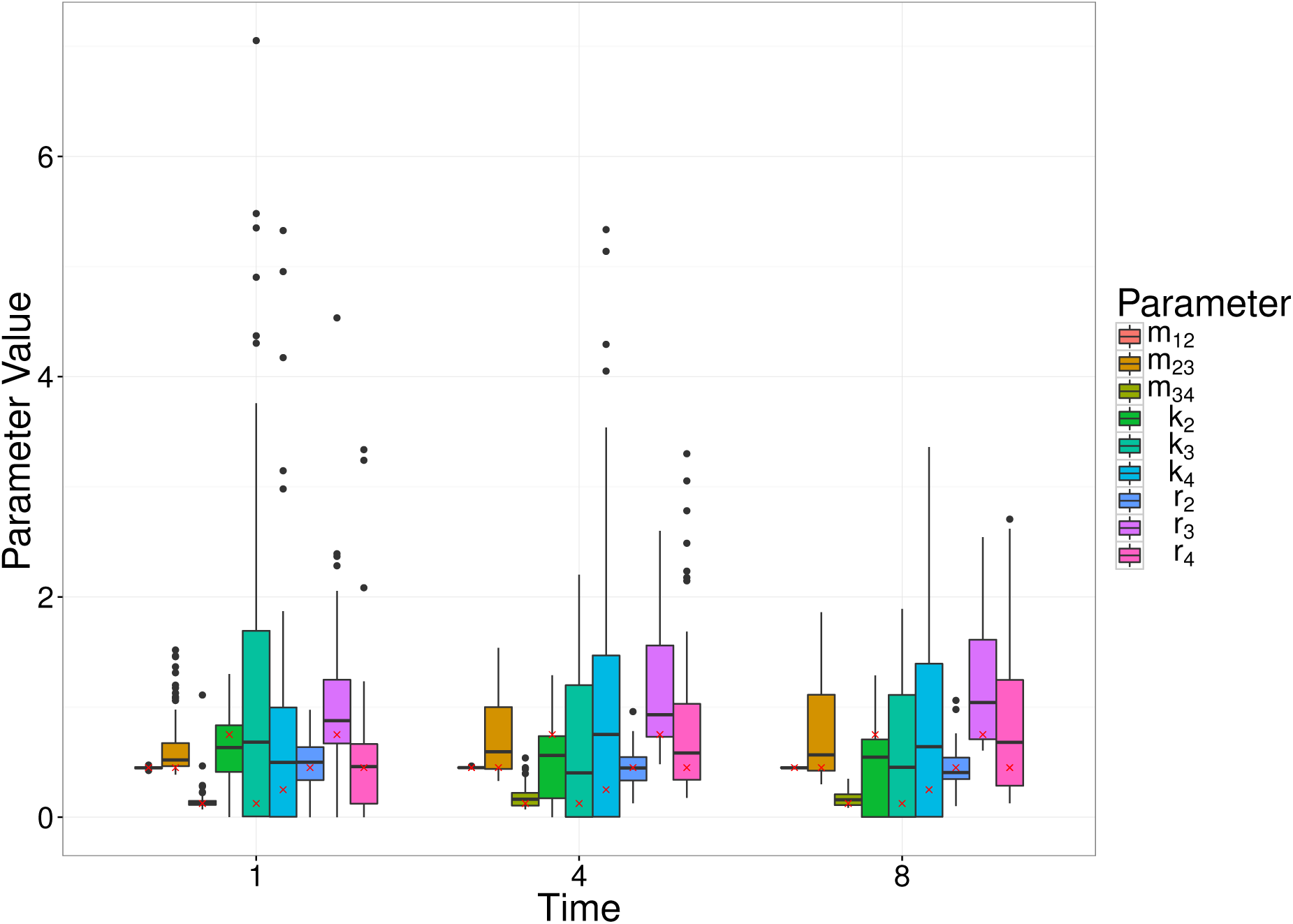
Box plots of MDE’s corresponding to simulated data for a random set of parameter values (Scenario 4), at a number of different observation times (1, 4 and 8) of the linear model. The box plots represent 100 parameter estimates corresponding to the 100 simulations, and the red crosses denote the true parameter value.

**Figure 8.**
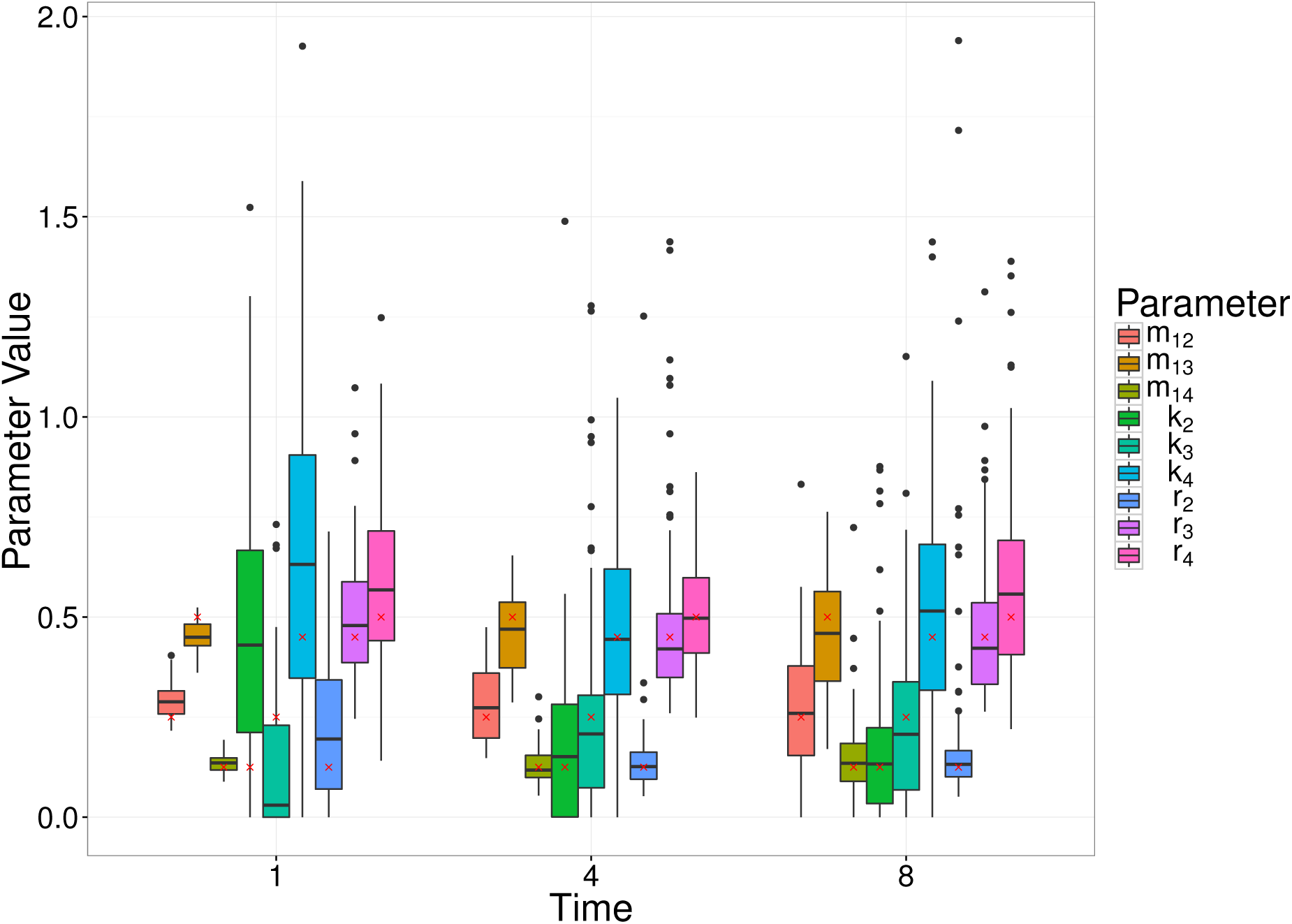
Box plots of MDE’s corresponding to simulated data for a random set of parameter values (Scenario 4), at a number of different observation times (1, 4 and 8) of the radial model. The box plots represent 100 parameter estimates corresponding to the 100 simulations, and the red crosses denote the true parameter value.

### 3.3 Migration in two directions

Many biological systems allow for migration in both directions between any two compartments in the system (e.g., later observations of the *Salmonella* system in [11] have shown that bacteria migrate back to the blood from the liver and spleen). Here, we consider a three compartment model where there is migration in both directions between two of the compartments, and we are trying to estimate the parameters, *m*_12_, *m*_13_, *m*_21_, *k*_2_, *k*_3_, *r*_2_, *r*_3_. Note that this example can be represented as either a radial or linear network. Similar to the migration birth-death model in previous examples, we consider all combinations of two representative parameter values (0.25, 0.50), and observing the process at times *t* = 1, 2, *…*, 8. Initial conditions are drawn from a Poisson distribution with mean parameter 200, and each experiment consists of 100 replicates (i.e., *A × T*). For each scenario (i.e., parameter combination and observation time), 100 experiments were simulated. As we established above that the Kullback-Leibler divergence was best, we only consider parameter estimation with respect to this divergence measure. The performance of the parameter inference is evaluated using the MARE, as before. Figure 9 shows the MARE across each observation time for the Kullback-Leibler divergence, and Figure 10 demonstrates the parameter estimates for three randomly selected scenarios. The MARE here is consistent with the Kullback-Leibler divergence estimates parameters in the previous example (Figure 4), noting the different number of parameters being estimated. Note in Figure 10, that even with a single observation time, the MDE method is able to identify all parameters.

**Figure 9.**
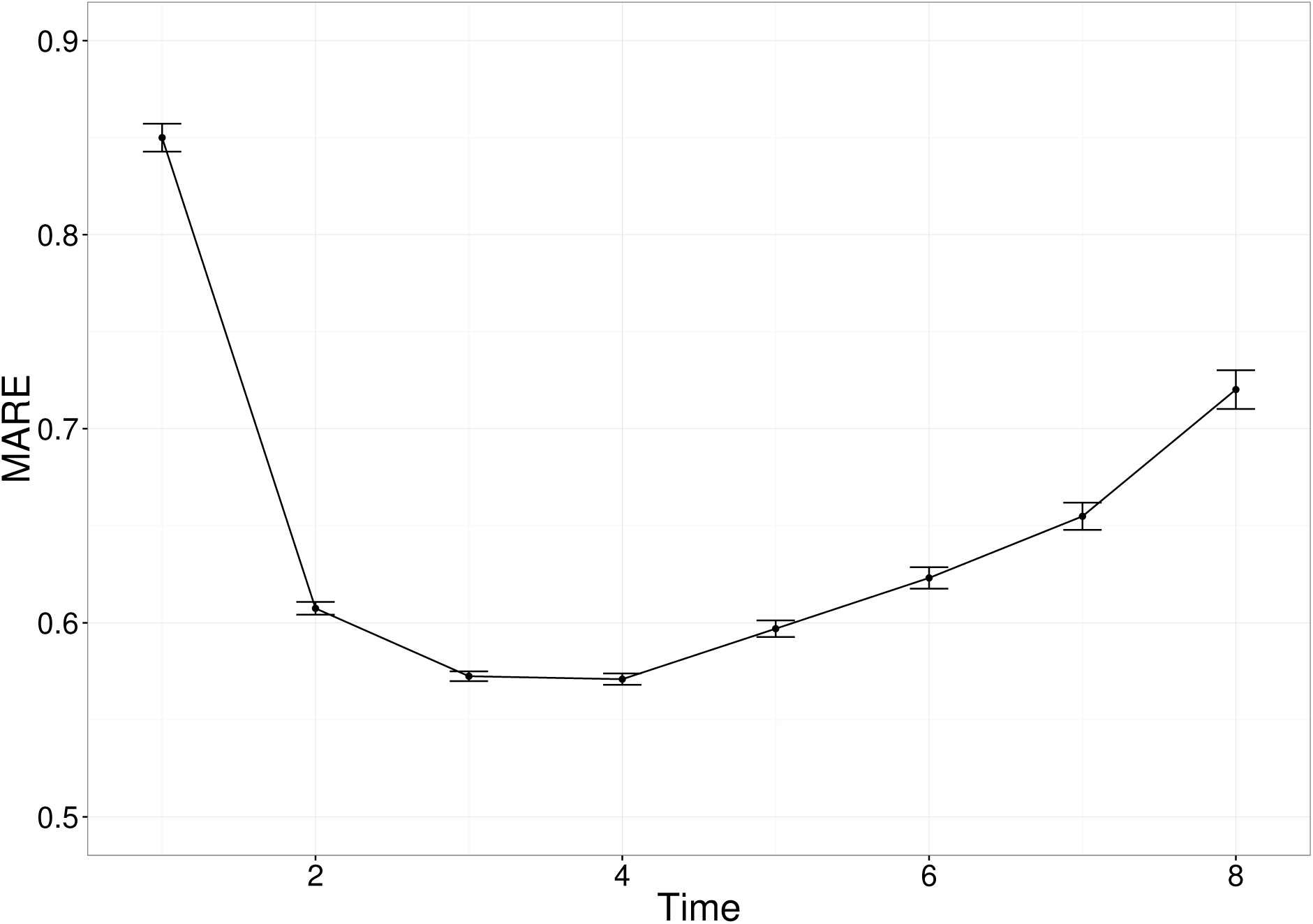
The average mean absolute relative error (MARE) across a number of observation times, for the three-compartment network with migration in both directions between two compartments. Each point is the average for that measure, at that observation time, with error bars representing the standard error.

**Figure 10.**
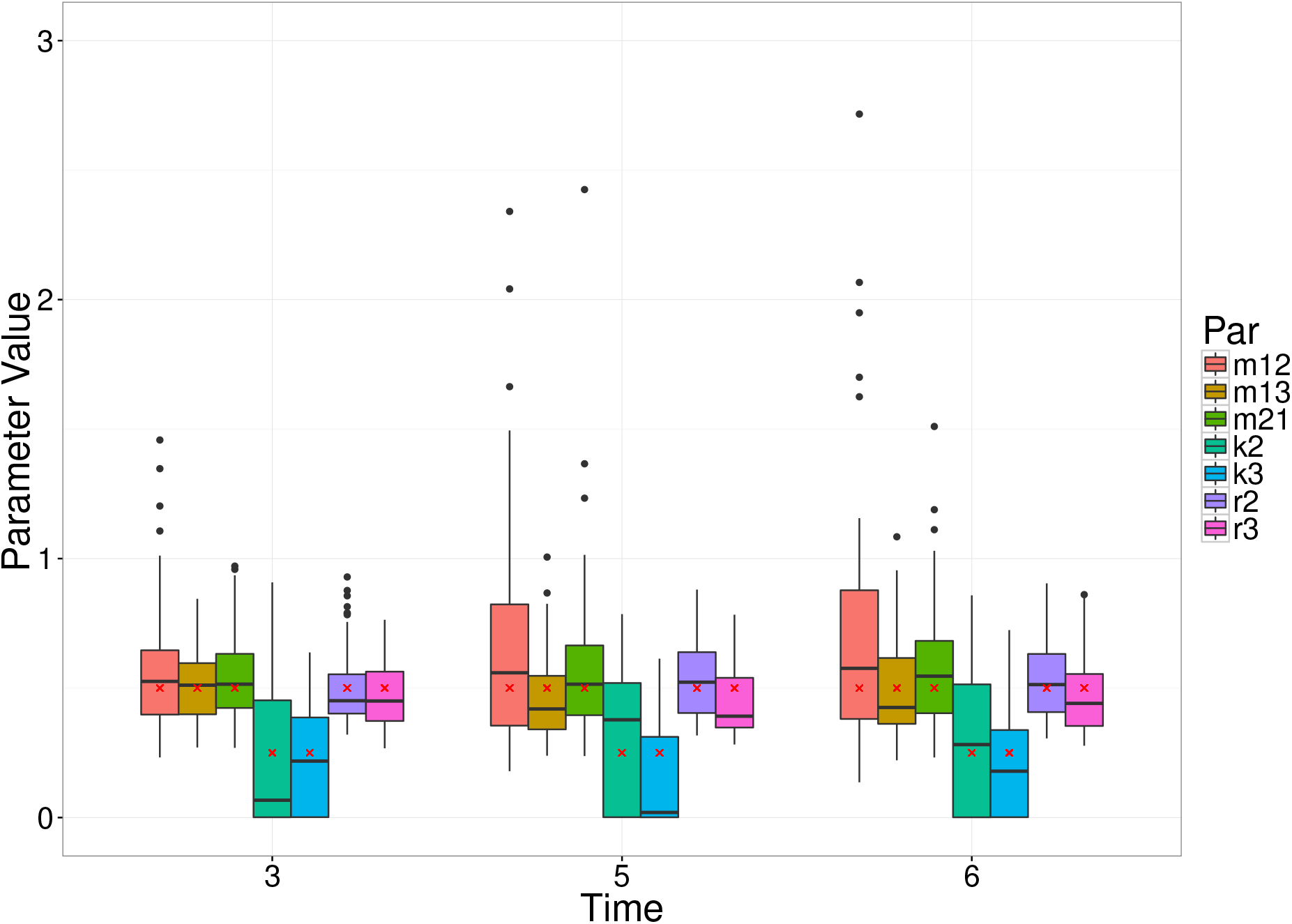
Box plots of MDE’s corresponding to simulated data for a random set of parameter values, at a number of different observation times of the three compartment model with migration in both directions between two compartments. The box plots represent 100 parameter estimates corresponding to the 100 simulations, and the red crosses denote the true parameter value.

### 3.4 Likelihood Comparison

In order to assess the overall quality of the inference achieved from this minimum divergence estimation technique, we compare the parameter inferences to those obtained via a Maximum Likelihood approach. The full details of the maximum likelihood estimation can be found in [11]. Briefly, a numerical procedure is used to solve the master equation for a given set of parameter values up to the required observation time. The probability of each observed value is then used to evaluate the likelihood of the observed data. An optimisation routine (UObyQA optimisation routine in the powell package [29], as above), is used to search across the parameter space for the maximum likelihood estimates. For simplicity, we once again consider the migration birth-death model (Figure 3a).

We consider different numbers of experimental replicates (*A × T* = 24, 40, 80), and a range of observation times (*t* = 0.5, 2, 4), with all combinations of three parameter values (0.25, 0.50, 0.75), for parameters *m*_12_, *k*_2_, *r*_2_. For each scenario (i.e., parameter values, experiment size, observation time), 30 simulations were produced, and initial conditions were randomly sampled from a Poisson distribution with mean parameter 50. We were not able to consider the same conditions as in previous examples, due to the computational burden of the likelihood-based method.

Figure 12 shows the difference in computation time between the moments-based method, and the likelihood-based method, across the different observation times. In particular, note that as we evaluate the moments exactly (using the matrix exponential calculation), the computation time is roughly constant across different observation times. However, the likelihood-based method requires solving a differential equation forward through time, and so the later the observation time, the longer it takes to obtain the parameter estimate. This highlights one of the computational bottlenecks regarding the use of a likelihood-based approach to analyse data of this type – particularly when considering later time points.

Figure 11 shows the MARE across the different scenarios, for the minimum divergence estimation method, compared to the likelihood-based method. In particular, we can see that the likelihood-based method achieves a lower accuracy than the moments approach. This is due to the numerical method used to approximate the likelihood [11]. In order to improve the efficiency, the probability mass of the system is evolved forward through time – considering only states corresponding to a probability above a predefined tolerance – and the relevant probabilities used. Reducing the tolerance would increase the accuracy of the likelihood-based method, but again, at a greater computational cost. As we can see in Figure 12, the likelihood-based method is already less computationally efficient, and to obtain a more accurate parameter estimate than the moments approach, would require significantly more time.

**Figure 11.**
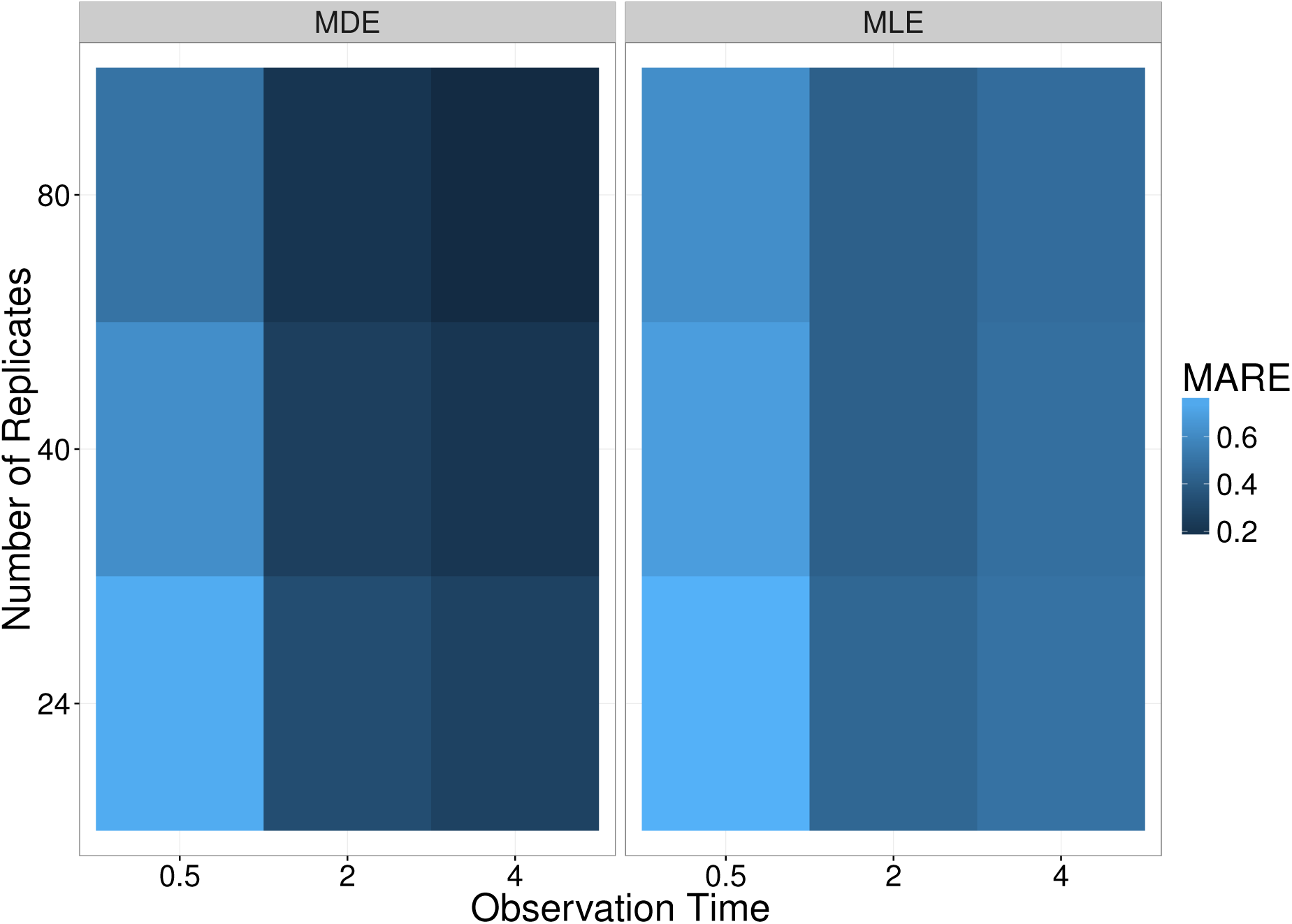
Comparison between MDE (left) and MLE (right) at inferring parameters of simulated data, for a range of different number of replicates (*A × T*), and different observation times. The colour of each panel represents the MARE across the 30 simulations, under the 27 different combinations of parameter values. The darker colour indicates less error in the estimates (on average) from simulate data. As one would expect, a larger number of tagged strains per experiment gives increased accuracy, and given the nature of this system (and the initial conditions), a later observation time corresponds to better accuracy – as the replication and killing rates are difficult to identify at early times.

**Figure 12.**
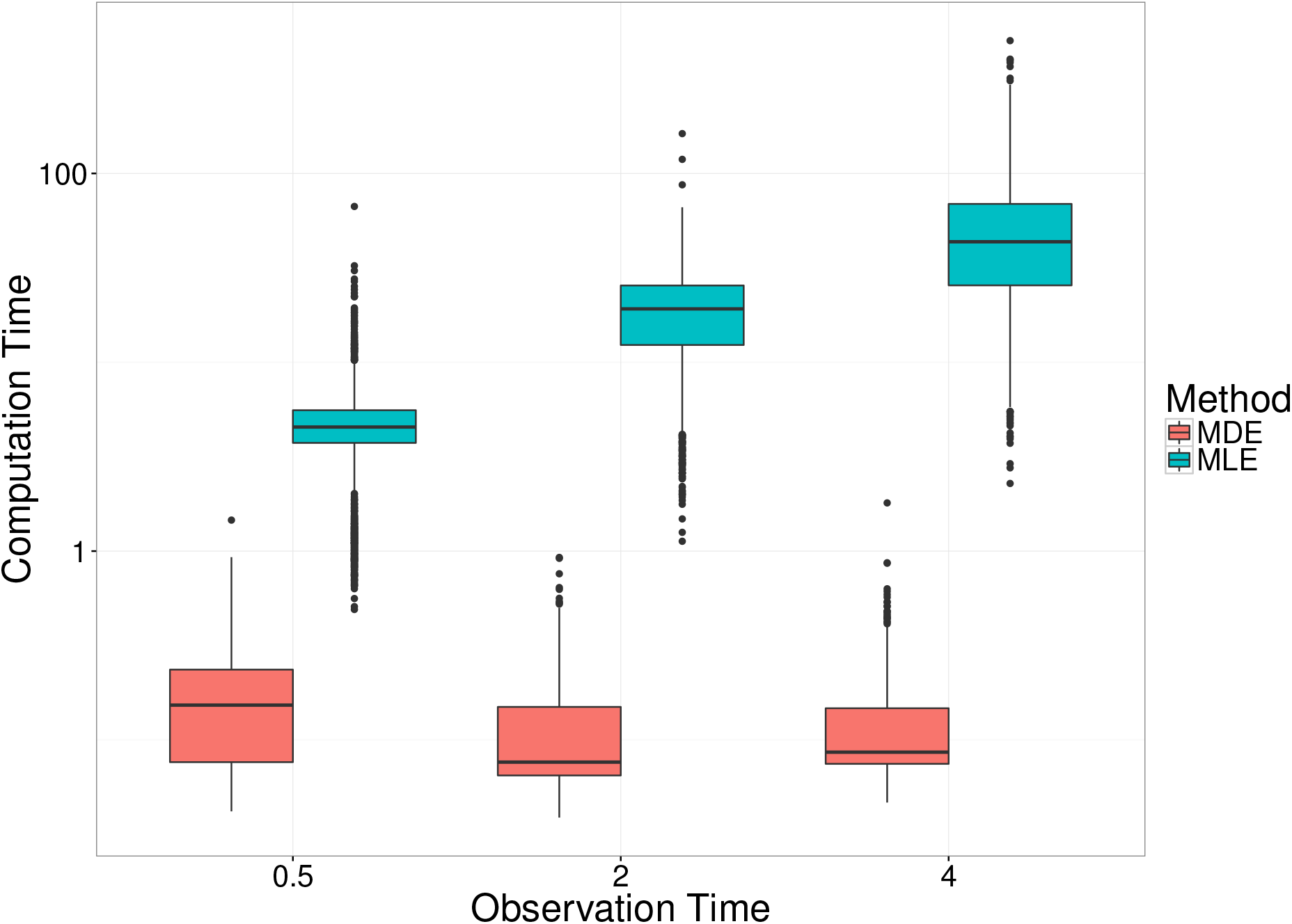
Comparison between MDE and MLE methods at inferring parameters of simulated data with respect to the computation time. Each box plot comprises of 30 simulations from 27 combinations of parameter values, for three different numbers of tagged strains. The size of the experiment does not impact the computation time for either method.

### 3.5 Analysis of experimental data

#### 3.5.1 Description of the system

Finally, we performed a re-analysis of experimental data from [11], in which groups of mice received an intravenous dose of *Salmonella enterica* Typhimurium, composed of an even mixture of 8 wildtype isogenic tagged strains (WITS). Bacterial loads and WITS composition were measured in the blood, liver and spleen of 10 mice at each observation time. During the first phase of the WITS experiments (represented by the first two time points at 0.5 and 6 hours post inoculation), the only biologically relevant processes considered are: migration of bacteria from blood to liver or spleen, replication and death inside the liver and the spleen. That is, on state space *S* = {(*n_B_, n_L_, n_S_*) *| n_B_, n_L_, n_S_ ≥* 0}, we have the following transition rates:

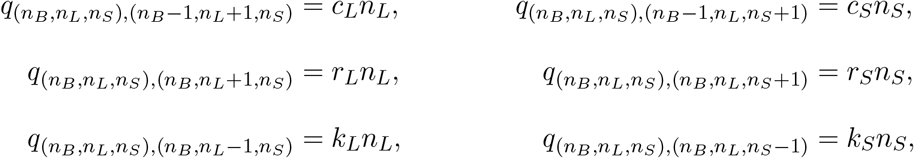

where *c_L_, c_S_* are the clearance (=migration) rates from the blood into the liver and spleen respectively, *r_L_, r_S_* are the replication rates in the liver and spleen respectively, and *k_L_, k_S_* are the killing rates in the liver and spleen, respectively.

One purpose of the study in [11] was to compare the effects of two vaccines on the dynamics of infection. Here, we re-analyse two of the experimental groups: an untreated group (naive) who received no vaccine, and a group who received a live-attenuated vaccine (LV). In the naive group, in addition to the clearance, replication and killing rates, we estimated the effective inoculum size *i*_0_ = *N_B_* (0) to account for the possibility that some of the inoculated bacteria failed to initiate an infection process [11]. In the vaccinated group, no bacteria were recovered from the blood at either of the two observation times, hence we were unable to estimate the corresponding migration rates. Instead we assumed that the transfer of bacteria to the organs was effectively instantaneous, and sought to estimate the effective initial loads in the liver and spleen (i.e., *i_L_* = *N_L_*(0) and *i_S_* = *N_S_* (0)), as well as the replication and killing rates. In each case, the inoculum size is assumed to be Poisson distributed.

#### 3.5.2 Parameter estimation with observation noise

We applied the MDE method to estimate the parameters corresponding to these systems, and compared the results to the estimates obtained via the MLE approach, presented in [11]. Having obtained the MDE parameter estimates, we used the bootstrap approach to calculate uncertainty intervals on the estimates, and assess the model goodness-of-fit.

As described in [11], the recorded data were subject to observation noise. In particular, only a fraction of some organs were sampled (e.g., as we are not able to fully recover the total amount of blood from an individual mouse), and there was noise introduced via the qPCR. The observed moments were adjusted using the simulation-based, pre-processing procedure described in Section 2.7: we assumed that the form of the sampling was binomial, and the qPCR noise has been previously modelled using a log-Normal distribution [11]. For example, for the naive group, we sampled 10,000 parameters (*c_L_, c_S_, k_L_, k_S_, r_L_, r_S_*) each independently from *U* [0, 3], and initial inoculum *i*_0_ *∼* Poisson(30.33) (the mean WITS population calculated from plating experiments [11]). “Perfect moments” were calculated from simulations of the same size as the experimental data. The corresponding observed moments were then calculated by applying the observation noise model – binomial sampling to represent the fractional sampling of organs within each mouse, and log-Normal noise to represent the qPCR noise – to each simulation, and calculating the moments. The variances were log-transformed for the purpose of the regression models to predict the corresponding perfect moments from those observed from the experiment.

Figure 13 provides a comparison of the MLE and MDE estimation procedures for the naive and LV experimental groups. The box plots represent the bootstrapped parameter estimates, with the MDE parameter estimate (blue dots), and MLE parameter estimates (red crosses). Figure 14 shows the estimated replication and killing rates in both the liver and spleen, for both groups. The cross and dashed ellipse (red) represent the MLE, and the dot and solid ellipse (blue), represent the MDE (where the ellipse is calculated from 1000 bootstrap samples, represented as grey dots). As explained in the previous section, the strong positive correlation between the replication and killing rates within each organ, clearly visible on Figure 14, is characteristic of this system.

**Figure 13.**
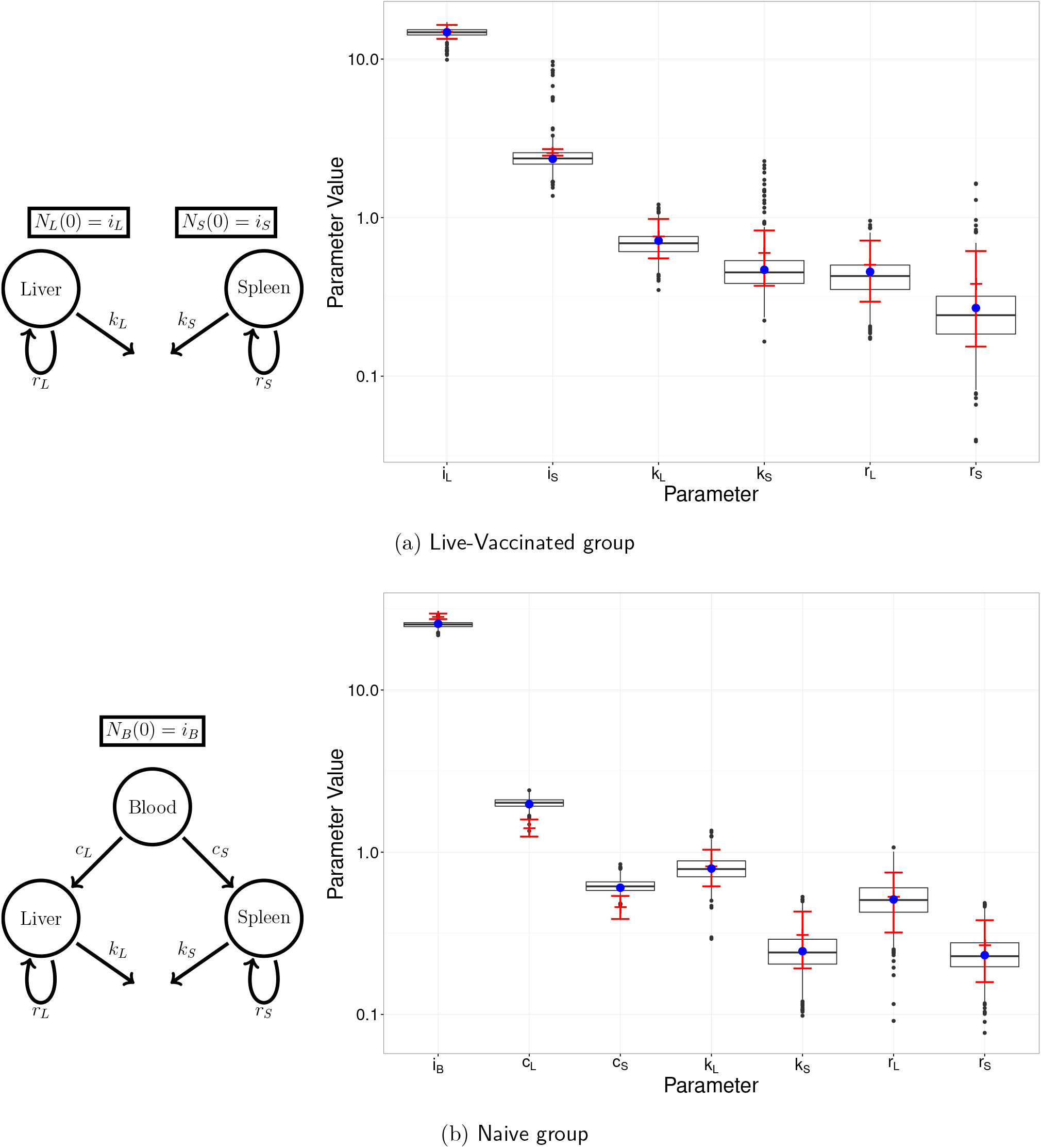
Left: Diagram of each model illustrating the relevant compartments, initial conditions, and rates of interest. Right: MDE parameter estimate (blue dot) with box plots of the bootstrapped parameter estimates, and MLE parameter estimate (red cross) with 95% confidence interval (red bars).

**Figure 14.**
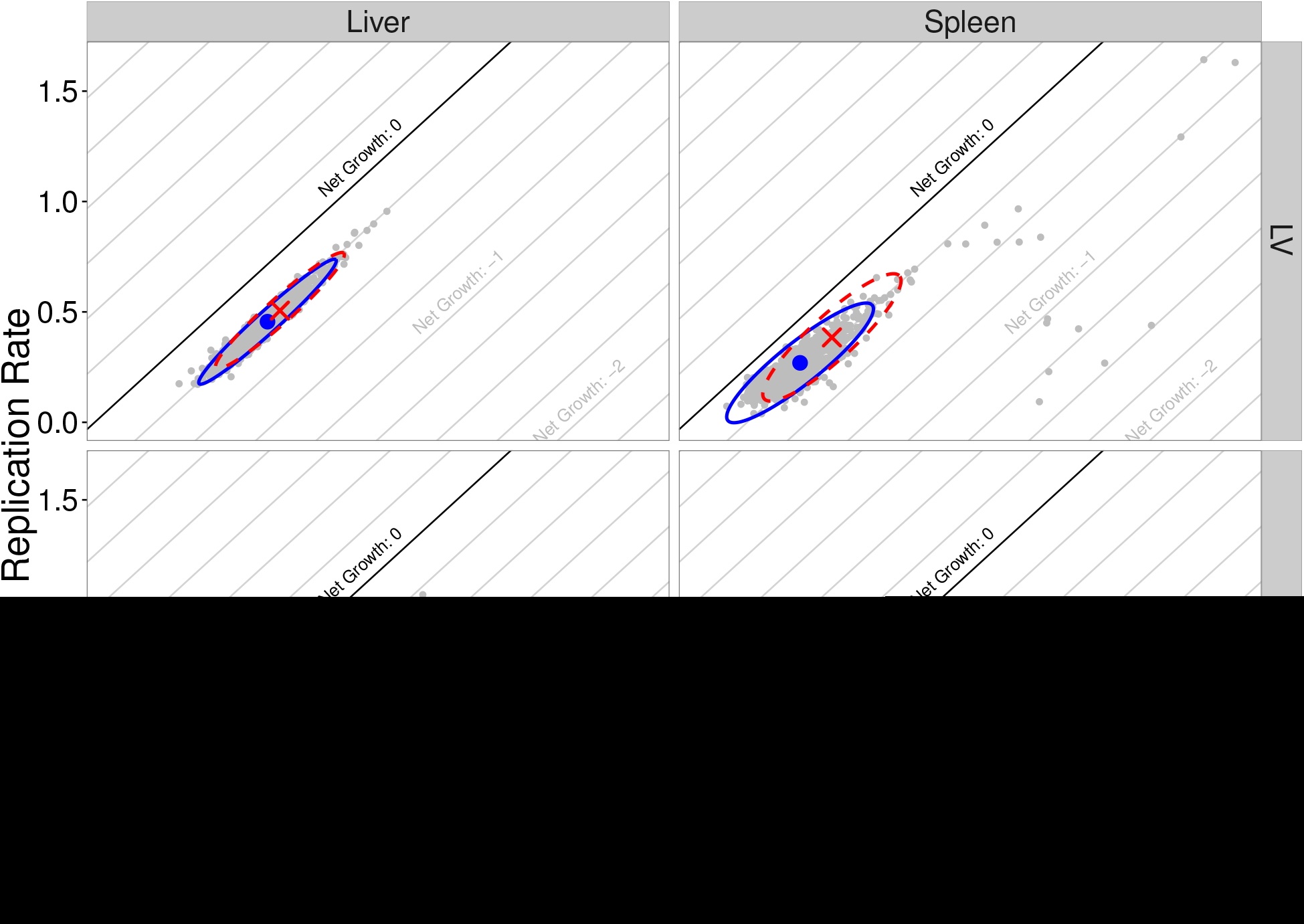
Bivariate distributions of replication and killing rates in liver and spleen, for both naive and live-vaccinated groups. Blue circles are the MDE values, with the blue (solid) ellipses representing the 95% confidence ellipses calculated using the 1000 bootstrap samples (grey points). The red crosses are the MLE values, with red (dashed) ellipse calculated using the hessian evaluated at the MLE.

#### 3.5.3 Model goodness-of-fit

In calculating the MDE parameter estimates for the bootstrap samples, we evaluated divergence measures for datasets simulated from the model. Thus, the collection of these divergences provide a suitable representation of the null distribution of divergences under this model. We compared the divergence for our observed data set to this null distribution in order to assess the model goodness-of-fit. Figure 15 demonstrates the goodness-of-fit measure for the models fit to the two experimental groups.

**Figure 15.**
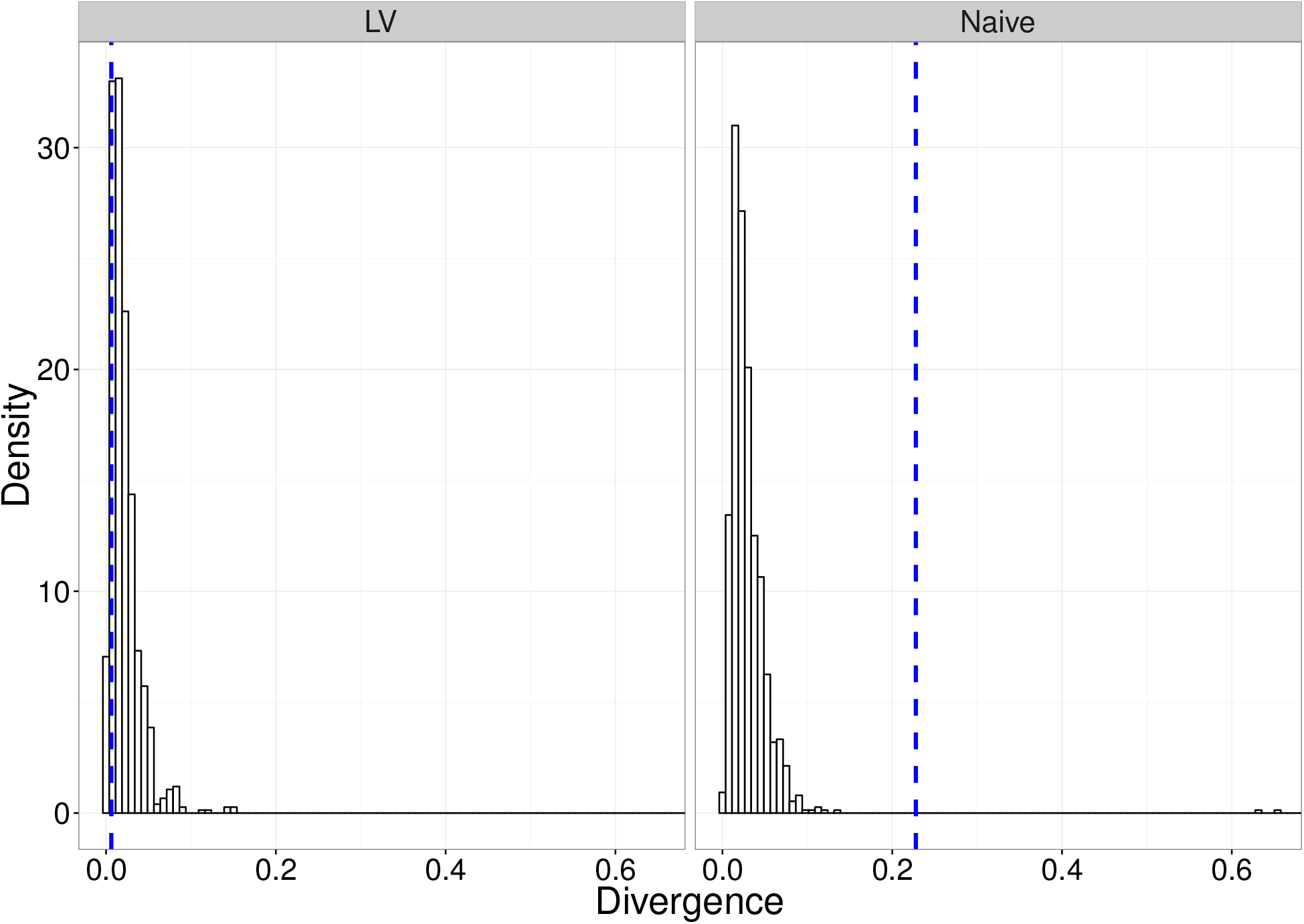
Figures demonstrating the goodness-of-fit of the two models to the respective data sets. The histogram bars are the bootstrapped estimate of the null distribution of divergences under the model at the estimated parameter values for the respective model. The (blue) vertical dashed-line is the divergence corresponding to the observed data set.

The approximate p-values associated with the goodness-of-fit tests for the naive and LV experimental groups are 0.002 and 0.882, respectively. This suggests that the model provides a suitable fit to the vaccinated experimental group; however, there is a non-negligible discrepancy between the model output for the naive group fit, and the observed data. This suggests that some of the assumptions in the original dynamic model may be erroneous; however, revisiting them goes beyond the scope of the present study.

As a further demonstration of the model goodness-of-fit, we compare the model output at each of the estimated parameter sets, to the observed data. Figure 16 shows these simulations from each of the MDE and MLE approaches, at each of the 0.5h and 6h observation times, for both the naive and vaccinated experimental groups. Observational noise consistent with that in the experimental data was added to the simulated data (i.e., binomial sampling for observed counts, and log-normal noise consistent with qPCR). The plots indicate that the main source of discrepancy between data and model in the naive-mice group is the distribution of bacterial loads in the blood after 6 hours.

**Figure 16.**
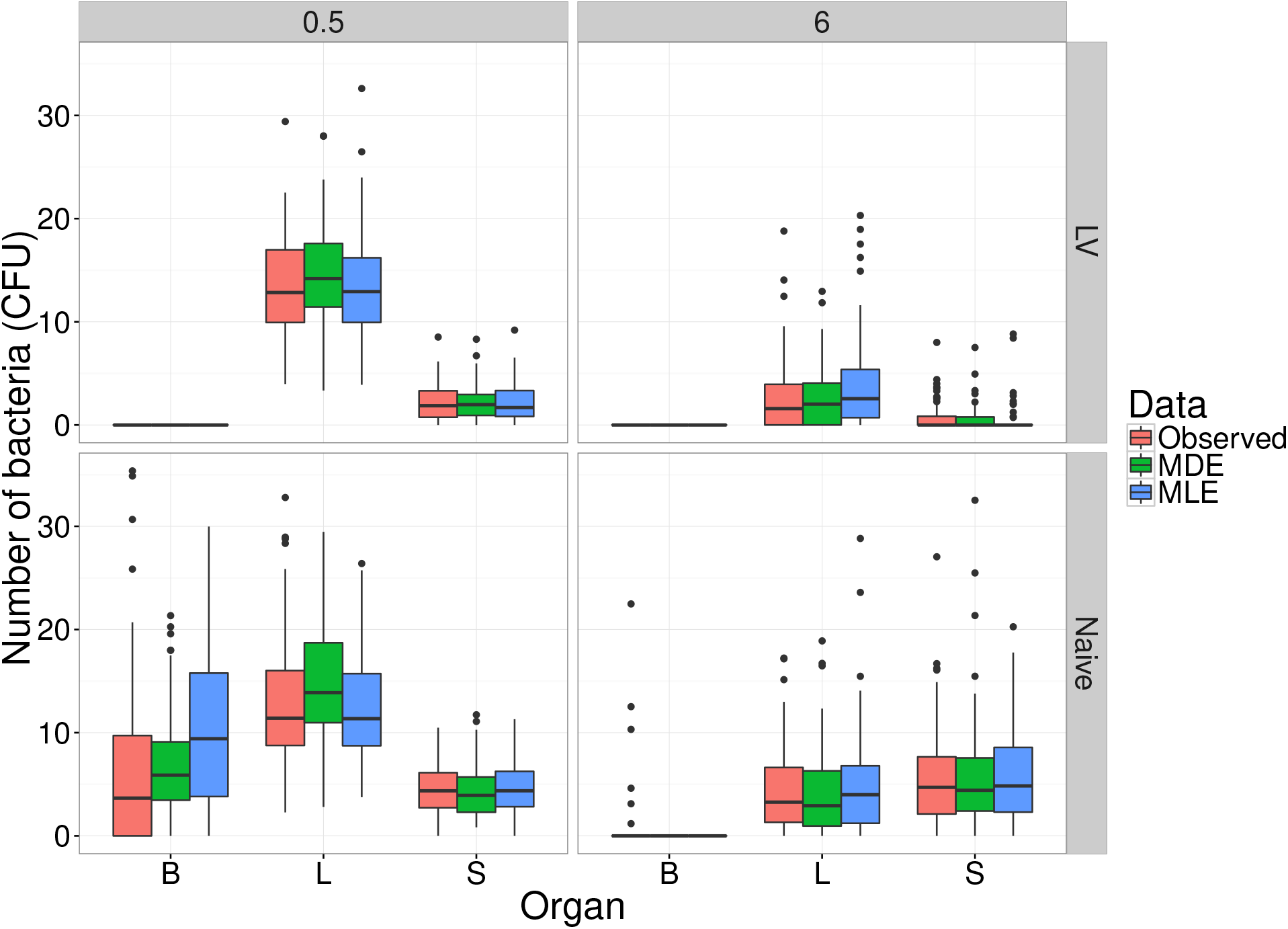
Observed experimental data for both naive and vaccinated groups, at both observation times, compared to simulations from the model at parameters estimated by both the MLE and MDE methods, in each of the blood (B), liver (L) and spleen (S). Numbers of bacteria are the sums of eight identical WITS.

## 4 Discussion

We have described a functional and flexible moments-based method to estimate parameters of stochastic metapopulation models, and applied it to experimental data on within-host bacterial infection dynamics. Compared to simulation-based methods, our technique delivers accurate estimation orders of magnitude faster. Although simulation-based inference has grown in popularity in computational biology—either within likelihood-free [17] or approximate-likelihood [16] approaches—even with the increasing availability of fast, multi-core computers, algorithms can still take days to converge in multi-parameter and multi-variable problems. While this is not a major issue when fitting a single model to a single dataset, it is a hindrance for more ambitious applications: indeed there is a growing demand in biology for model selection and model-based optimal experimental design, which are much more computationally intensive tasks (e.g., [34, 35, 36, 37]).

In its current form, our method can incorporate any linear, multivariate, continuous-time Markovian model, and fit it to experimental data that include multiple replicates and an arbitrary number of independent time points. These last two characteristics are typical of experimental biology, yet little attention has been given to systems of this form in recent statistical developments. Indeed, many inference methods for stochastic models (for example, driven by applications to epidemiology), target inference from a single time-series [38, 16].

In addition, we have provided a worked example for how to correct for observation noise, applicable to any given experimental system. While sampling error is typically taken into account in likelihood-based inference, many inference studies either ignore experimental noise (e.g. [24]) or choose arbitrary distributions (e.g. [39]). Even if experimental error has limited impact on the mean of the observations, it will affect the variance, with implications for the precision and reliability of statistical inference. The method we demonstrated here is based on pre-processing experimental data to effect an empirical correction of the observation noise. Crucially, it is not limited by mathematical tractability when combining multiple error sources, and it does not slow down the inference computation (i.e., it only needs to be performed once before fitting the model, and the same simulations are also used to adjust the moments of the noisy-simulated data).

The choice of a metric to minimise for likelihood-free inference is a common issue in computational statistics, especially in the context of increasingly popular ABC methods [40]. When the data and the model output are summarised by multiple statistics, the default option is to use either euclidian distance or a chi-squared type variant. The latter was chosen, for example, in a recent study using moment-based inference to solve a systems-biology problem [24]. However, because the summary statistics of interest are the moments of statistical distributions, we hypothesised it would be more informative to use a divergence metric instead. While many divergence measures exist for the purpose of comparing mathematically defined distributions [41], we are not aware of standard methods to compute divergence between multivariate distributions generated by complex stochastic processes. Instead, we took a pragmatic approach and used mathematical expressions for the Hellinger distance or the Kullback-Liebler divergence between multivariate normal distributions, because these expressions depend on the first-and second-order moments of the distributions only. Even though the resulting divergence measures are not the actual Hellinger distance or Kullback-Liebler divergence between our observed and predicted distributions, they still outperform the chi-squared metric based on parameter inference accuracy. It is worth noting that, across the three model structures we tested, there was not a single metric that was consistently better than any other, and the differences in accuracy were often relatively small (Fig. 4). Depending on the degree of accuracy sought, it may be worth testing these and other metrics against simulations before applying this method to a different experimental system. It is our intention to provide a flexible blueprint that can be tailored to other problems, rather than a one-size-fits-all black box which may prove unreliable as soon as the circumstances change.

The particular molecular technology (isogenic tagging) that motivated the development of this inference method, has become pervasive in the study of within-host dynamics of bacterial infection in the last 10 years (see reviews by [42, 2, 43]). Yet, to our knowledge, this is the first attempt to provide a general modelling and inference framework that could be applied to any of these experimental systems. Indeed, previous efforts have been tailored to specific case studies [43, 7, 9, 13, 44, 8] despite asking fundamentally similar questions: how fast are bacteria replicating and dying? How much migration is taking place among organs or tissues? As soon as any two of these dynamic processes are co-occurring, it is not possible to evaluate them based solely on average bacterial loads: it is necessary to obtain reliable estimates of the variance, preferably within a single animal, by quantifying a set of independent and isogenic tags. The first known example goes back 60 years, using two naturally occurring mutants of *S. enterica* that could be distinguished by selective growth medium to investigate colonisation dynamics in mice. Although a wide range of bacterial tagging methods (including antibiotic markers and fluorophores) have been used since, non-coding DNA barcodes have opened new prospects as arbitrarily large numbers of truly isogenic tags can be generated and quantified by sequencing [14]. It is our hope that the inference framework presented here will contribute to the field’s extension by providing much needed analytical support to analyse and design microbiology experiments.

## Acknowledgements

We thank the many microbiologists whose collaboration has helped initiate and shape this study, including Andrew Grant, Eric Harvill, Emma Slack, Chris Coward and Duncan Maskell. We are also grateful to fellow modellers Roland Regoes, Simon Frost, Julia Gog and Joshua Ross for their technical advice.

